# RNA helicase, DDX27 regulates skeletal muscle growth and regeneration by modulation of translational processes

**DOI:** 10.1101/125484

**Authors:** Alexis H Bennett, Marie-Francoise O’Donohue, Stacey R. Gundry, Aye T. Chan, Jeffery Widrick, Isabelle Draper, Anirban Chakraborty, Yi Zhou, Leonard I. Zon, Pierre-Emmanuel Gleizes, Alan H. Beggs, Vandana A Gupta

**Author notes:** Corresponding Author, Vandana A Gupta, Division of Genetics, Department of Medicine, Brigham and Women’s Hospital, Harvard Medical School, Boston, MA 02115.

## Abstract

Gene expression in a tissue-specific context depends on the combined efforts of epigenetic, transcriptional and post-transcriptional processes that lead to the production of specific proteins that are important determinants of cellular identity. Ribosomes are a central component of the protein biosynthesis machinery in cells; however, their regulatory roles in the translational control of gene expression in skeletal muscle remain to be defined. In a genetic screen to identify critical regulators of myogenesis, we identified a DEAD-Box RNA helicase, DDX27, that is required for skeletal muscle growth and regeneration. We demonstrate that DDX27 regulates ribosomal RNA (rRNA) maturation, and thereby the ribosome biogenesis and the translation of specific transcripts during myogenesis. These findings provide insight into the translational regulation of gene expression in myogenesis and suggest novel functions for ribosomes in regulating gene expression in skeletal muscles.

**AUTHOR SUMMARY:** Inherited skeletal muscle diseases are the most common form of genetic disorders with primary abnormalities in the structure and function of skeletal muscle resulting in the impaired locomotion in affected patients. A major hindrance to the development of effective therapies is a lack of understanding of biological processes that promote skeletal muscle growth. By performing a forward genetic screen in zebrafish we have identified mutation in a RNA helicase that leads to perturbations of ribosomal biogenesis pathway and impairs skeletal muscle growth and regeneration. Therefore, our studies have identified novel ribosome-based disease processes that may be therapeutic modulated to restore muscle function in skeletal muscle diseases.

## INTRODUCTION

Ribosome biogenesis is fundamental to all life forms and is the primary determinant of translational capacity of the cell. Ribosome biogenesis is a complex process that involves transcription, modification and processing of ribosomal RNA, production of ribosomal proteins and auxiliary factors and coordinated assembly of ribonucleoprotein complexes to produce functional ribosomes [1]. While previously considered a “house keeping” constitutive process, recent studies have shown that ribosome biogenesis is regulated differently between cells and can be modulated in a cell type-specific manner [2]. These differences are required to generate ribosomes of different heterogeneities and functionalities that contribute to the translational control of gene regulation by selecting mRNA subsets to be translated under specific growth conditions potentially by identifying specific recognition elements in the mRNA [3-5]. Protein synthesis is the end stage of the gene regulation hierarchy and despite the identification of translational regulators of specific genes, a systematic identification of translational regulatory processes critical for tissue specific control of gene expression is still lacking. Moreover, upstream regulatory factors/processes that regulate ribosome heterogeneity in a cellular and organ-specific context in vertebrates still remains to be identified.

Skeletal muscle is a contractile, post-mitotic tissue that accounts for 30-50% of body mass. Skeletal muscle growth and repair is a highly coordinated process that is dependent on the proliferative expansion and differentiation of muscle stem cells that originate from the muscle precursor cells at the end of embryogenesis [6-8]. Activated muscle stem cells through symmetric/ asymmetric divisions give rise to daughter cells that maintain the muscle stem cell population (satellite cells) and committed myogenic progenitor cells (MPC) that fuse to form the differentiated skeletal muscle [6, 9, 10]. Defects in processes regulating muscle stem cells, myoblast fusion and differentiation therefore, constitute pathological pathways affecting the muscle growth and regeneration [11-17]. Although epigenetic and transcriptional controls of myogenesis have been studied extensively, the importance of translational regulation of these processes in skeletal muscle function is less defined. Transcriptional and translational analysis of myoblasts has shown that differential mRNA translation controls protein expression of specific subset of genes during myogenesis. Recent studies have revealed ribosomal changes during skeletal muscle growth and atrophy. An increase in ribosome biogenesis is often observed during skeletal muscle hypertrophy [18, 19]. Ribosomal perturbations on the other hand are associated with skeletal muscle growth and diseases [20-23]. Deletion of ribosomal protein genes *S6k1* and *Rps6* in mice results in smaller myofibers and reduced muscle function. In several muscular dystrophy models and atrophied muscles, a reduction in ribosome number and/or activity is observed [22, 24, 25]. In spite of these dynamic changes observed in ribosomes during different growth conditions, our understanding of the mechanism(s) that regulate ribosomal biogenesis in skeletal muscle and eventually control the translational landscape during muscle growth is still poor.

In a forward genetic screen to identify critical regulators of myogenesis *in vivo*, we recently identified a RNA helicase gene, *ddx27*, that controls skeletal muscle growth and regeneration in zebrafish [26, 27]. We further show that DDX27 is required for rRNA maturation and ribosome biogenesis in skeletal muscles. Strikingly, DDX27-deficient myoblasts exhibit impaired translation of mRNA transcripts that have been shown to control proliferation as well as differentiation of muscle progenitors during myogenesis. These studies highlight the specific role of upstream ribosome biogenesis processes in regulating tissue specific gene expression during myogenesis.

## RESULTS

### Identification of novel regulators of myogenesis *in vivo*

Zebrafish and human exhibit similar skeletal muscle structure and the molecular regulatory hierarchy of myogenesis is conserved between zebrafish and mammals [9, 28]. Therefore, to identify regulators of skeletal muscle growth and disease *in vivo*, we performed an ENU mutagenesis screen in zebrafish [26]. Analysis of skeletal muscle of 4-5 dpf (days post fertilization) larvae of one mutant identified in the screen, *Osoi* (Japanese for “slow”), displayed highly reduced birefringence in polarized light microscopy in comparison to the control, indicative of skeletal muscle defects (Fig 1A). Positional mapping and sequencing of this mutation identified a 20 bp deletion in exon 18 of the *ddx27* gene (DEAD-box containing RNA helicase 27) (Fig 1B, S1A and B). qRT-PCR analysis showed significantly lower levels of *ddx27* transcripts reflecting probable nonsense-mediated decay (S1C Fig). Overexpression of human *DDX27* resulted in a rescue of skeletal muscle defects in mutant embryos, demonstrating functional evolutionary conservation and confirming that mutation in *ddx27* are causal for the *osoi* phenotype (Fig 1C). Evaluation of a large number of mutant embryos showed that homozygotes die by 6-7 dpf. qRT-PCR analysis of mouse *Ddx27* mRNA demonstrated *Ddx27* expression in several tissue types (*S*1D Fig). Whole-mount immunofluorescence of wild-type zebrafish (4 dpf) detected Ddx27 expression in Pax7 labeled muscle progenitor cells in skeletal muscle (Fig 1D).

**Figure 1.**
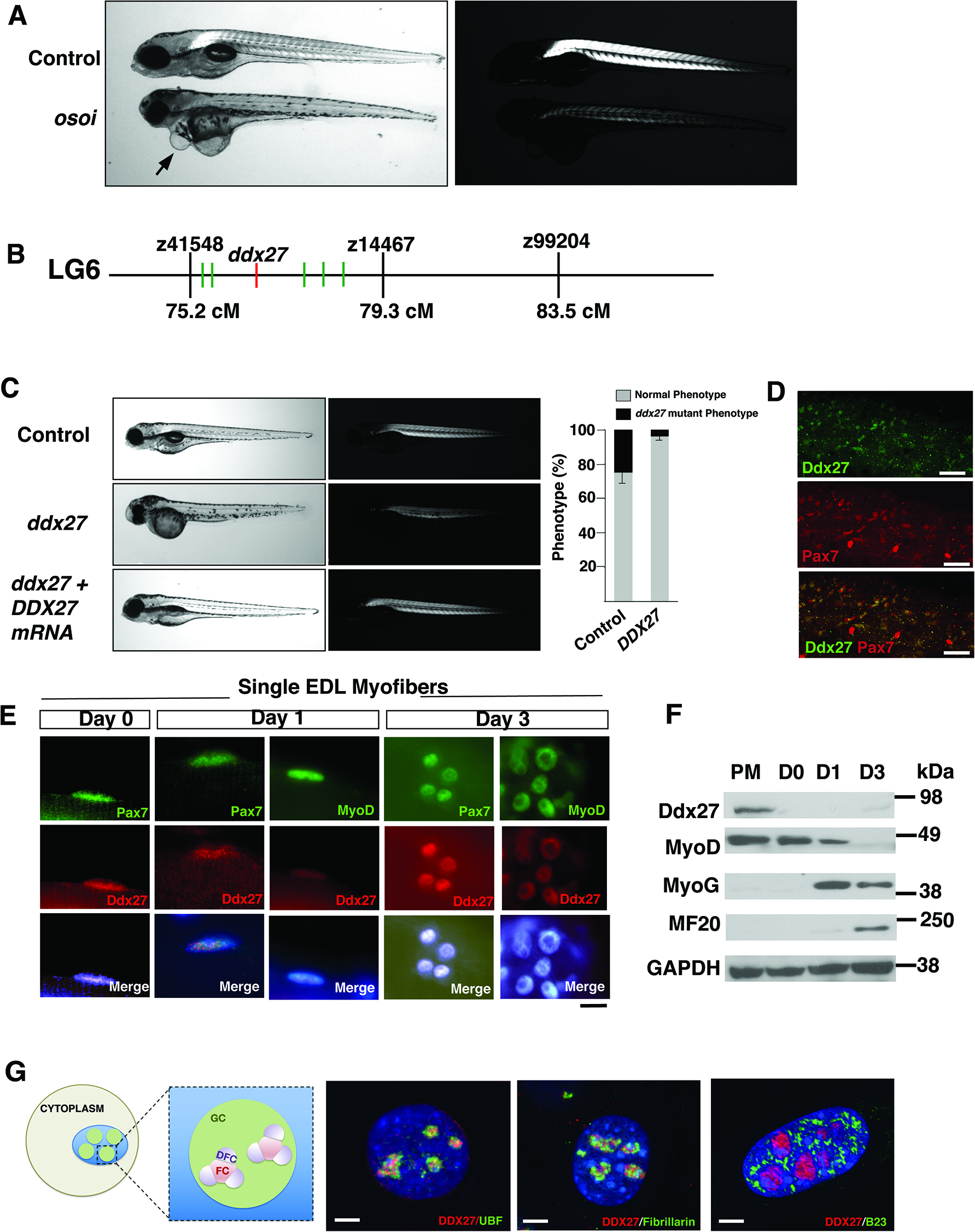
Mutation in a nucleolar protein *ddx27* results in skeletal muscle abnormalities. (A) Microscopic visualization of control and mutant larval zebrafish (*osoi*) at 5 days post fertilization (dpf). Mutant fish display leaner muscles (left panel) and exhibit highly reduced birefringence in comparison to control (right panel). Mutant fish also exhibit pericardial edema (arrow) (B) Genetic mapping of *osoi* mutant by initial bulk segregant analysis identified linkage on chromosome 6. Fine mapping of chromosome 6 resolved flanking markers z41548 and z14467, with a candidate genome region containing six candidate genes that were sequenced by Sanger sequencing (C) Overexpression of human *DDX27* mRNA results in a significant decrease in mutant zebrafish phenotype (D) Whole-mount Immunofluorescence was performed on control and *ddx27* mutant larvae (4dpf) (scale bar: 50μm) (E) Immunofluorescence on newly isolated (Day 0) and cultured (Day1 and 2) EDL myofibers from wild-type mice (scale bar: 10μm). (F) Western blot showing relative expression of Ddx27 and myogenic markers (MyoD, MyoG and MF20) in proliferating C2C12 myoblasts in growth media (50% confluence) or in differentiation media for 3 days (D0-3). GAPDH was used as the control. (G) Schematic diagram of nucleolus depicting nucleolar domains. Eukaryotic nucleolus has tripartite architecture: Fibrillar center (FC); Dense fibrillar component (DFC) and granular compartment (GC). Immunofluorescence of human myoblasts with DDX27 and nucleolar markers labeling each compartment of nucleolus (scale bar: 2μm)

To identify DDX27 expression domains in mammalian skeletal muscle, immunofluorescence was performed on single myofibers isolated from extensor digitorum longus (EDL) muscles of wild-type mice (Fig 1E). Immunofluorescence on freshly isolated EDL myofibers (Day 0) showed that Ddx27 is expressed in Pax7 positive satellite cells. EDL myofibers were cultured for 3 days, which results in activation (day1), proliferation and differentiation of satellite cells (day 2). Ddx27 expression remained high in activated satellite cells (day1). Ddx27 expression was observed in proliferating Pax7 positive nuclei as well as MyoD expressing nuclei. Subsequently, after culture of myofibers for 1-2 days, satellite cells typically undergo cell division. After 3 days in culture, proliferating satellite cells as well as proliferating myoblasts (MyoD positive) exhibited Ddx27 expression. To determine Ddx27 expression during muscle differentiation, western blot was performed on proliferating and differentiating C2C12 myoblasts (Fig 1F). Ddx27 expression was detected in proliferating myoblasts and declined as cells are committed towards differentiation (100% confluence, day 0). The downregulation of ddx27 corresponded to an increase in MyoG and myosin heavy chain expression (MF20). These data show that Ddx27 expression is high in proliferating satellite cells and myoblasts and is reduced in myoblasts as they are committed towards the terminal differentiation into myotubes.

Finally, analysis of subnuclear localization by immunofluorescence revealed that DDX27 is co-localized with UBF (fibrillar component) and Fibrillarin (dense fibrillary component) in nucleoli of human myoblasts (Fig 1G). No co-localization of ddx27 was detected with B23, labeling the granular component of nucleoli. The nucleolus is the primary site of ribosome biogenesis, therefore, this sub-nucleolar organization of DDX27 suggests a functional requirement of DDX27 in rDNA transcription and/or pre-rRNA processing steps in skeletal muscles.

### Precocious skeletal muscle differentiation in Ddx27 deficiency

To investigate the role of RNA helicase Ddx27 in skeletal muscle homeostasis, we analyzed the structure and function of mutant zebrafish embryos (2 dpf) and larvae (4-5 dpf. Analysis of skeletal muscle histology at the end of primary myogenesis in 2 dpf embryos showed no visible differences between control and mutant skeletal muscles (S2A Fig). Quantification of fiber cross section area (CSA) also showed no significant differences in control and mutant muscles at 2 dpf (S2B Fig). The nuclear content of the myofibers was analyzed as fiber CSA per nuclei that was similar in control and mutant myofibers at 2 dpf, however showed a significant decrease in mutants at 4 dpf (S2C Fig). This suggests that skeletal muscle development is normal during embryogenesis (0-2 dpf) in *ddx27* fish. During development, Ddx27 expression is observed during embryogenesis and persists during larval stages in zebrafish (www.zfin.com). However, normal skeletal muscle growth during embryogenesis could be due to a functional redundancy with other family members, many of which have overlapping expression patterns with Ddx27. Homozygous *ddx27* mutant fish (5 dpf) showed reduced fiber diameter (Fig 2A-B arrow, inset) and central nuclei as observed in several muscle diseases (Fig 2A-B, arrowhead) [29]. Electron microscopy of longitudinal sections of *ddx27* mutant myofibers also showed disorganized myofibrillar structures in *ddx27* mutant fish (Fig 2C-E, arrow). The mutant myofibers displayed smaller Z-lines and disorganized actin-myosin assemblies in comparison to controls suggesting differentiation defects of these muscles. Immunofluorescence of cultured myofibers from control and mutant zebrafish further showed a reduction and disorganization of skeletal muscle differentiation markers at 4 dpf, (Actn2/3 and Ryr1) validating our earlier observation that absence of Ddx27 results in a defective differentiation of skeletal muscles (S2B Fig). The centralized mutant nuclei were round in shape with highly enlarged nucleoli in comparison to controls (Fig 2E, arrowhead). Cross-sections of muscles showed a reduction in myofiber diameter (52 ± 14%) in *ddx27* mutant fish in comparison to controls (Fig 2F-H). Skeletal muscles of mutant fish also displayed whorled membrane structures associated with nuclei (Fig 2D, arrow). These whorled membrane structures are often observed in skeletal muscle of congenital myopathy patients, however their origin and biological significance remains unknown. Ddx27 belongs to a family of highly conserved RNA helicases in vertebrates. Therefore, to evaluate if the function of Ddx27 in skeletal muscle is conserved in vertebrates, myogenic differentiation of control and *Ddx27* knockout C2C12 myoblasts was analyzed. C2C12 myoblasts exhibited reduced proliferation and impaired differentiation, that failed to form mature myofibers (S2F-G Fig). These results show that Ddx27 deficiency affects skeletal muscle growth during zebrafish larval stages leading in to skeletal muscle hypotrophy and DDX27 functions in growth and differentiation of skeletal muscle are conserved among vertebrates.

**Figure 2.**
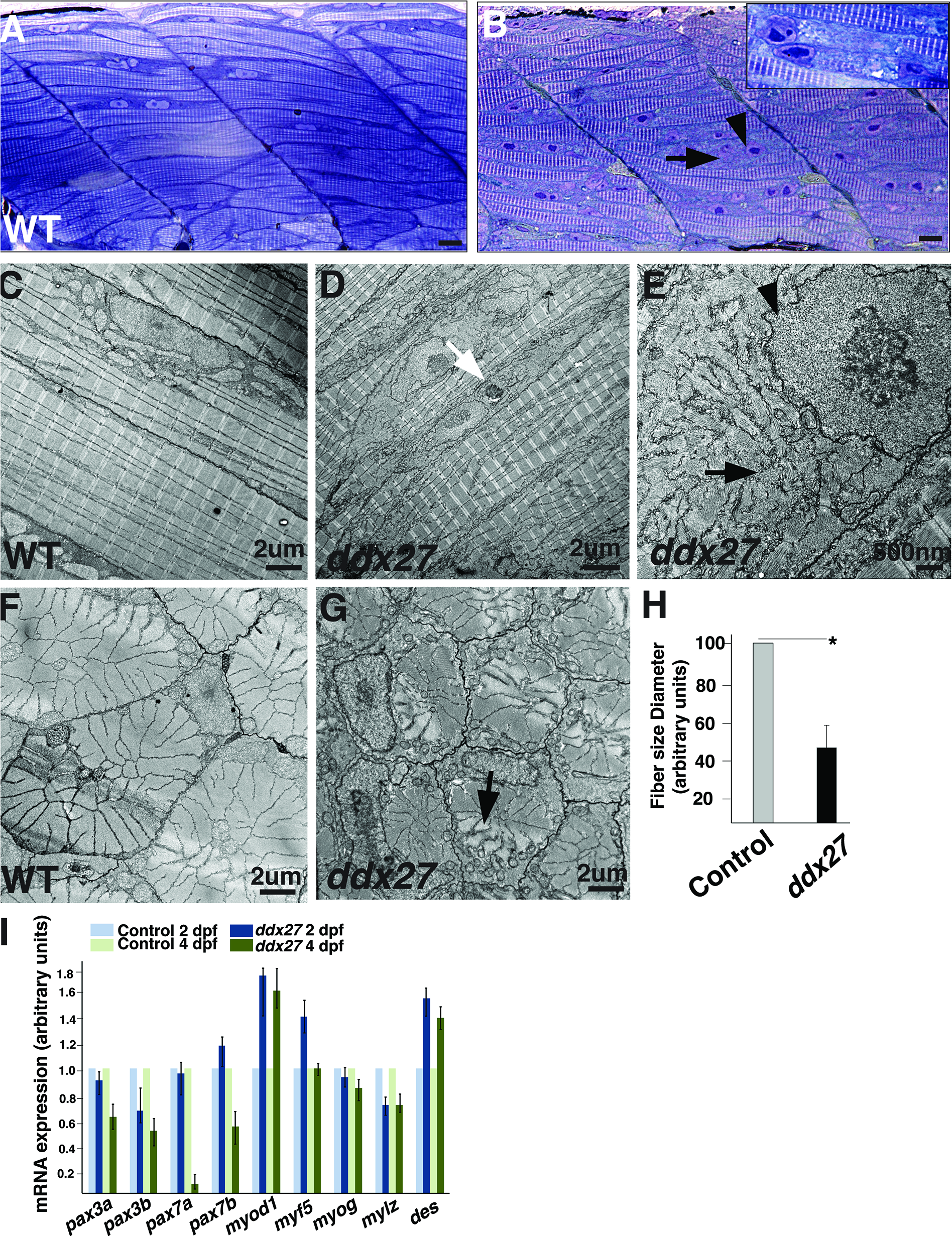
Ddx27 deficiency results in skeletal muscle hypotrophy and precocious differentiation. (A-B) Histology of longitudinal skeletal muscle sections in control and *ddx27* mutant stained with toluidine blue exhibiting enlarged nuclei (arrowhead) and areas lacking sarcomeres (arrow) at 5 dpf. High magnification view (boxed area) (C-G) Transmission electron micrographs of skeletal muscles (longitudinal view: C-E, and cross-section view: F-G) in control and *ddx27* mutant (5 dpf). Arrows indicating disorganized sarcomere (H) Quantification of myofiber size in control and *ddx27* mutant fish (5 dpf) (n=10) (I) qRT-PCR of control and *ddx27* mutant fish showed a reduction in the expression of muscle stem cell markers (*pax3, pax7*) and an increase in expression of myogenic commitment genes (*myod1* and *myf5*). The expression of late differentiation markers was reduced in *ddx27* fish suggesting that pre-maturation expression of early myogenic genes results in abnormal disorganization of skeletal muscles.

To identify molecular events leading to growth defects in *ddx27* mutant zebrafish, qRT-PCR was performed during embryonic (2 dpf) and larval (4 dpf) stages. qRT-PCR revealed no significant expression changes in muscle progenitor cell markers, *pax3* and *pax7* during the embryonic stage (Fig 2I). In zebrafish, the *PAX7* orthologue is duplicated in two copies; *pax7a* and *pax7b. pax7a* expressing cells act to initiate myofiber formation post-injury, whereas *pax7a/b* expressing cells are required for myofiber growth [30]. Significant down regulation of both *pax7a and b* was observed during the larval stage in *ddx27* mutant zebrafish. Concurrently, with a downregulation of *pax3* and *pax7a/b*, we also observed a high expression of *myoD, myf5* and *desmin* in mutant muscles suggesting a premature activation of the myogenic program in Ddx27 deficiency (Fig 2I). However, this precocious activation of the myogenic program appears to be unable to progress to a proper differentiated state, as documented by low levels of late differentiation markers *myog* and *mylz* leading to disorganized sarcomeres.

### Reduced contractile strength in Ddx27-deficient skeletal muscles

*ddx27* zebrafish larvae exhibit significantly slower swimming than wild-type controls at 5 dpf (91 ± 29 mm/10 min for *ddx27* vs.16823 ± 214 mm/10 min for controls). To quantify the functional deficits in skeletal muscles of homozygous *ddx27* mutant larvae (5 dpf), peak twitch and tetanic forces of individual zebrafish skeletal muscle preparations were measured following electrical stimulation. Substantially depressed twitch and tetanic forces as well as the slower rise and fall of tension were observed in *ddx27* mutants (Fig 3A). Absolute tetanic force was significantly less for *ddx27* larvae compared to controls (mean ± SD: 0.28 ± 0.18 vs. 1.43 ± 0.27 mN; p < 0.0001) (Fig 3B). These force deficits in *ddx27* persisted in the force measurements normalized for cross-sectional areas (Fig 3C: 11 ± 8 vs. 48 ± 6 kPa; p < 0.0001), suggesting that the depressed tetanic force of the *ddx27* mutant preparations is primarily due to intrinsic skeletal muscle deficits. Mutant *ddx27* preparations also showed significant reductions in mean absolute twitch force (Fig 3D: 0.16 ± 0.12 vs. 1.11 ± 0.23 mN; p < 0.0001) and twitch force normalized to the CSA of individual larvae (Fig 3E: 7 ± 5 vs. 37 ± 4 kPa; p < 0.0001). Interestingly, twitch force was depressed to a relatively greater extent than tetanic force as revealed by the significantly reduced twitch to tetanic force ratios of the *ddx27* preparations (Fig 3F: 0.57 ± 0.15 vs. 0.78 ± 0.06 kPa; p = 0.0006). In addition to these differences in force magnitude, the kinetics of force development and relaxation were also affected in mutant skeletal muscles. The *ddx27* larvae displayed a significantly slower rise in the maximal rate of twitch tension development, +dP/dt (Fig 3G: 1.06 ± 0.71 vs. 8.00 ± 1.30 kPa/ms; p < 0.0001), and a significantly slower maximal rate of twitch tension relaxation, −dP/dt (Fig. 3H: −0.47 ± 0.24 vs. −2.55 ± 0.35 kPa/ms; p < 0.0001). This disproportionate reduction in twitch vs. tetanic force and the slowing of twitch kinetics indicate a reduction in the quality of the contractile performance of the *ddx27* larvae that are consistent with reduced motility of mutant muscles.

**Figure 3.**
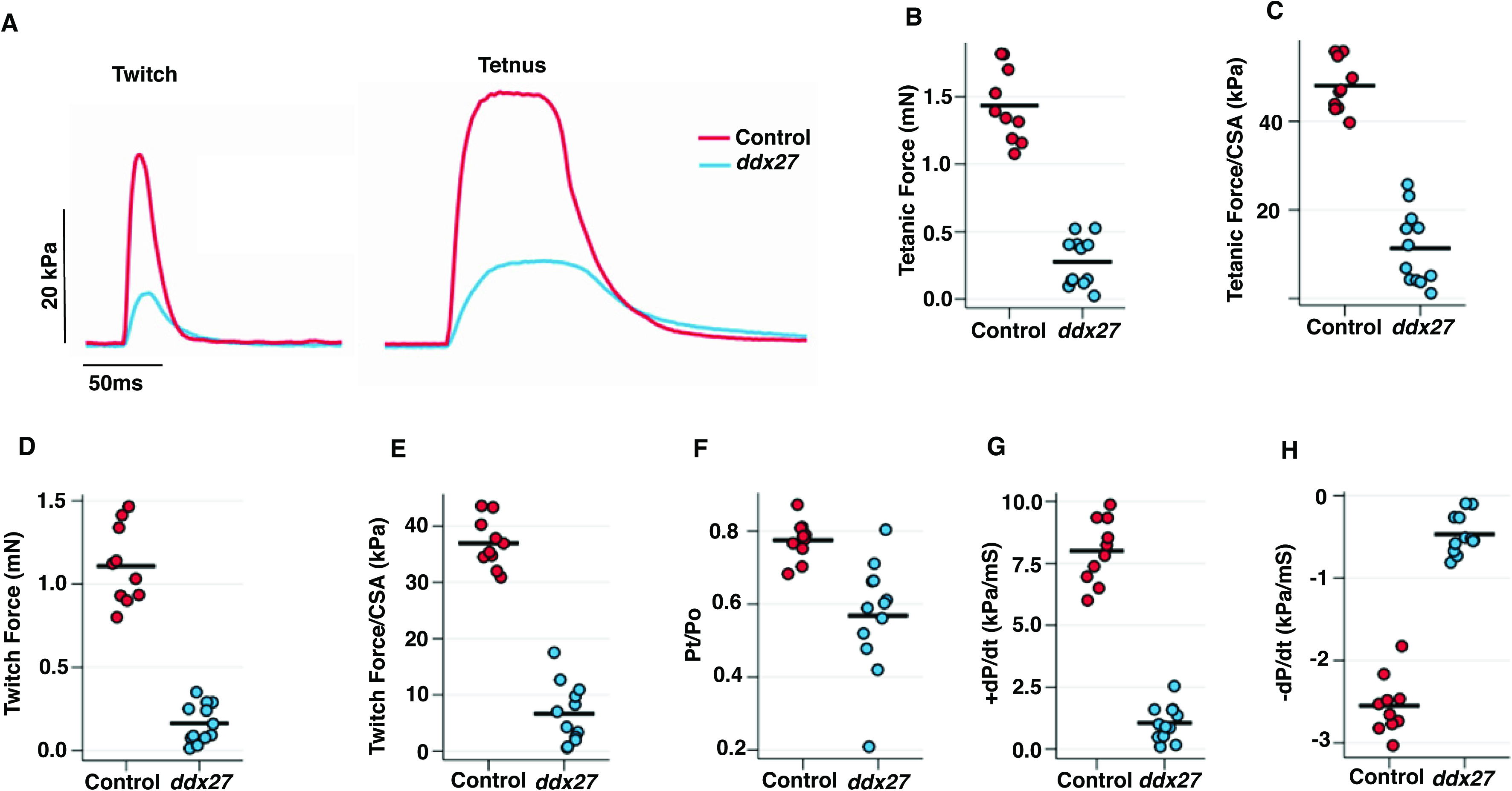
Ddx27-deficient skeletal muscle produce decreased contractile force and prolonged muscle relaxation. (A) Representative twitch (left) and tetanic force (right) records from control and *ddx26* zebrafish (5 dpf) preparation (B) Peak tetanic force (C) Peak tetanic force normalized to preparation cross-sectional area. Tetanic force is significantly reduced in *ddx27* mutant fish (D) Peak twitch force (E) Peak twitch force normalized to preparation cross-sectional area. Twitch force is significantly reduced in *ddx27* mutant fish (F) Maximal rate of twitch tension development. (G) Maximum rate of twitch force relaxation. (H) Twitch to tetanic force ratio. Each symbol represents an individual control (n = 10) or *ddx27* (n = 12) preparation with the group mean indicated by a horizontal line. t-tests indicated significant differences between control and mutant means for all 7 variables (P < 0.001 to P < 0.0001). Abbreviations: CSA, cross-sectional area; +dP/dt, maximal rate of tension development; −dP/dt, maximal rate of tension relaxation; Pt, peak twitch force; Po, peak tetanic force.

### Reduced Pax7-positive cell proliferation and impaired regeneration in Ddx27 deficiency

To understand if the decreased Pax7 expression observed in *ddx27* mutant is due to reduced Pax7 expression in MPCs or is caused by reduced number of MPCs, we performed whole mount immunofluorescence with Pax7 antibody during embryonic and larval stages. In zebrafish, Pax7 labels proliferative cells in dermomyotome, quiescent muscle stem cells and myoblasts that are required for muscle growth and regeneration [8, 31]. Quantification of Pax7 positive cells by whole mount immunofluorescence showed no differences in control and mutant muscles during embryogenesis (2 dpf). However, a significant reduction in Pax7 expressing nuclei (40 ± 8.2%) was observed in larval mutant skeletal muscle compared with control fish (4 dpf) (Fig 4A). This implies that reduced *pax7* expression observed in *ddx27* mutant fish is a consequence of reduced number of Pax7 cells in these fish. This decrease in Pax7 cells in mutants could either be due to a defect in proliferation of MPC population or an increased apoptosis in Ddx27 deficiency. To study the proliferative potential of Ddx27-deficient MPC, we performed 5-ethynyl-2’-deoxyuridine (EdU) incorporation assay and Pax7 immunofluorescence in control and mutant fish. The proportion of nuclei that were double labeled with Pax7^+^Edu^+^ in total Pax7 labeled population, decreased significantly to ~14.1% in *ddx27* skeletal muscle in comparison to the control (~22.0%). This indicates that the proliferation of Pax7 cells is significantly reduced in *ddx27* mutant fish (Fig 4B). We also investigated if Ddx27 deficiency leads to increased apoptosis of Pax7 MPC population. Whole mount TUNEL staining and western blot analysis with caspase 3 antibody at 3 and 4 dpf in control and mutant fish did not reveal any significant increase in apoptotic changes in mutant muscles compared to controls at these stages suggesting that decreased proliferation associated with premature differentiation rather than enhanced cell death underlie reduced MPC population observed in mutant muscles (S3 Fig).

**Figure 4.**
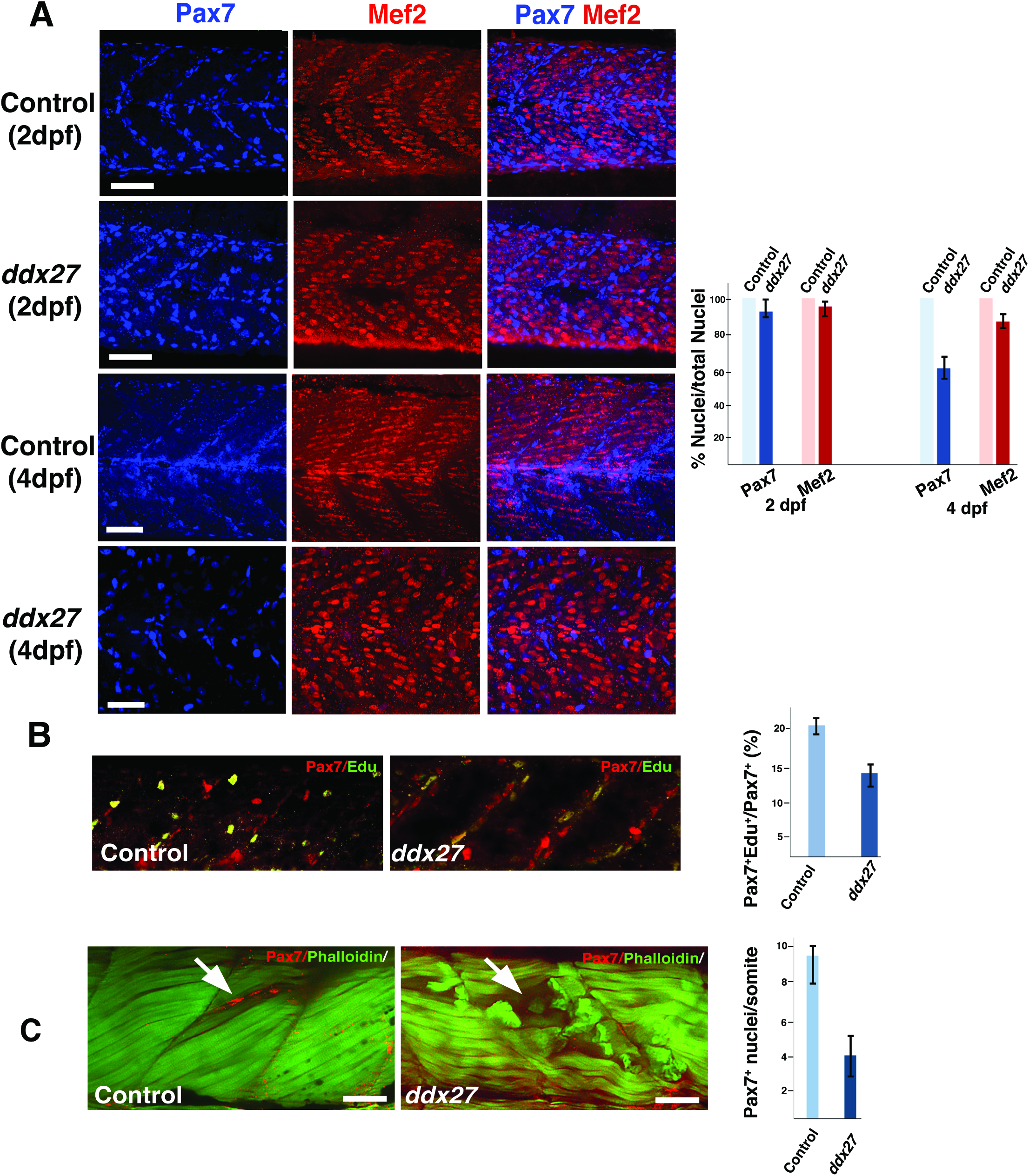
Ddx27 deficiency results in a decrease in muscle precursor cells (MPC) proliferation and skeletal muscle regeneration. (A) Whole mount immunofluorescence of zebrafish at different time intervals (2 dpf and 4 dpf) with MPC marker (Pax7) and late differentiation marker (Mef2) demonstrating a decrease in of MPC in *ddx27* mutant during post-embryonic skeletal muscle growth (4 dpf) (scale bar: 50μm) (B) Control and *ddx27* mutant zebrafish were pulse-labeled with EdU for 2hr and immunostained with Pax7. Fish were analyzed for EdU and Pax7 labeling (4 dpf) by whole mount immunofluorescence. The proportion of proliferative Pax7 population was estimated by quantifying Pax7^+^/Edu^+^ double-positive nuclei out of total Pax7^+^ nuclei in control and mutant fish (scale bar: 50μm) (C) Trunk muscles in control and *ddx27* zebrafish were injected with cardiotoxin (3 dpf). Skeletal muscles were analyzed at 5 dpf by whole mount immunofluorescence with Pax7 and phalloidin. Control muscles show an accumulation of Pax7 expressing cells at the site of injury (arrow) that was lacking in *ddx27* muscles (arrow) (scale bar: 50μm)

A stem cell niche equivalent to mammalian satellite cell system exists in zebrafish that involves migration and asymmetric division and/or proliferation of Pax7 expressing muscle progenitor cells to repair muscle upon injury [8, 32, 33]. Moreover, processes and timing of skeletal muscle repair in larval zebrafish are highly similar to adult mammalian skeletal muscle. As Pax7 MPC population also contributes to skeletal muscle repair in zebrafish, we next investigated if Ddx27 deficiency affects muscle repair in mutant fish. To induce muscle injury, cardiotoxin was injected into the epiaxial myotome of somites of larval fish at 3 dpf and skeletal muscles were analyzed at 5 dpf as described previously [9]. Whole mount immunofluorescence showed that cardiotoxin administration resulted in accumulation of a pool of Pax7 expressing cells at the site of injury in control fish which exhibited efficient myofiber repair post injury as seen by newly formed lighter stained myosin myofibers (Fig 4C). In contrast, injured Ddx27-deficient zebrafish muscles exhibited numerous degenerating myofibers and no accumulation of Pax7 expressing cells or muscle repair was observed post injury suggesting an impaired regeneration (Fig 4C). These findings suggest that Ddx27 plays a pivotal role in skeletal muscle regeneration. The major steps in skeletal muscle repair in zebrafish involve proliferation of MPC population, migration to the injury site and fusion with the damaged myofibers or form new myofibers [9]. *ddx27* mutant fish exhibit reduced MPCs proliferation as well as differentiation defects to form mature myofibers suggesting that either or both of these processes could underlie the repair defects observed in Ddx27 deficiency in skeletal muscle.

### Nucleolar abnormalities and rRNA maturation defects in Ddx27 deficiency

Ddx27 is localized in the nucleolus that is primarily the site of ribosome biogenesis. Therefore, we evaluated nucleolar structure and functions to understand the impact of Ddx27 on these processes in skeletal muscle. Immunofluorescence with different nucleolar markers showed changes in localization of the fibrillary component marker Ubf (labeling rRNA transcription sites) from small punctate foci to larger condensed areas, suggesting a perturbation in active transcription sites in the nucleolus. Similarly, fibrillarin-enriched dense fibrillary component areas of early rRNA-processing regions were also disrupted and merged, forming larger, more condensed structures. Lastly, granular component of the nucleolus which is the site of late rRNA processing was also perturbed in the *ddx27* mutant. B23, a nucleolar granular component marker exhibited altered localization to nucleoplasm in mutant nucleoli in comparison to the nucleolar restricted expression controls suggesting a structural disruption of fibrillary and dense fibrillary component potentially disrupts the organization of granular compartment in mutant nuclei (Fig. 5A). These results suggest that Ddx27 deficiency disrupts sites of rRNA synthesis, processing and early ribosomal assembly.

To investigate the effect of Ddx27 on rRNA synthesis, we performed *in situ* rRNA transcription analysis in zebrafish skeletal muscles (5 dpf) by 5-EU labeling of newly synthesized rRNA. Following 5-EU labeling, immunofluorescence was performed to visualize MPCs (Pax7 positive cells) or myonuclei (Actn2/3 labeling). Analysis of 5-EU signal in MPC population revealed a significant reduction in rRNA transcripts in Pax7 positive cells in *ddx27* fish (30 ± 8 %). Interestingly, a significant decrease in rRNA synthesis was also observed in myonuclei in control myofibers. Further, examination of mutant myonuclei revealed a reduction in rRNA synthesis in comparison to control myonuclei. Together, these results suggest that Ddx27 deficiency impairs rRNA synthesis directly in MPCs. As these MPCs differentiation to form myofibers, rRNA synthesis defect persists in myonuclei contributing to impaired ribosome biogenesis and reduced muscle function (Fig 5B).

**Figure 5.**
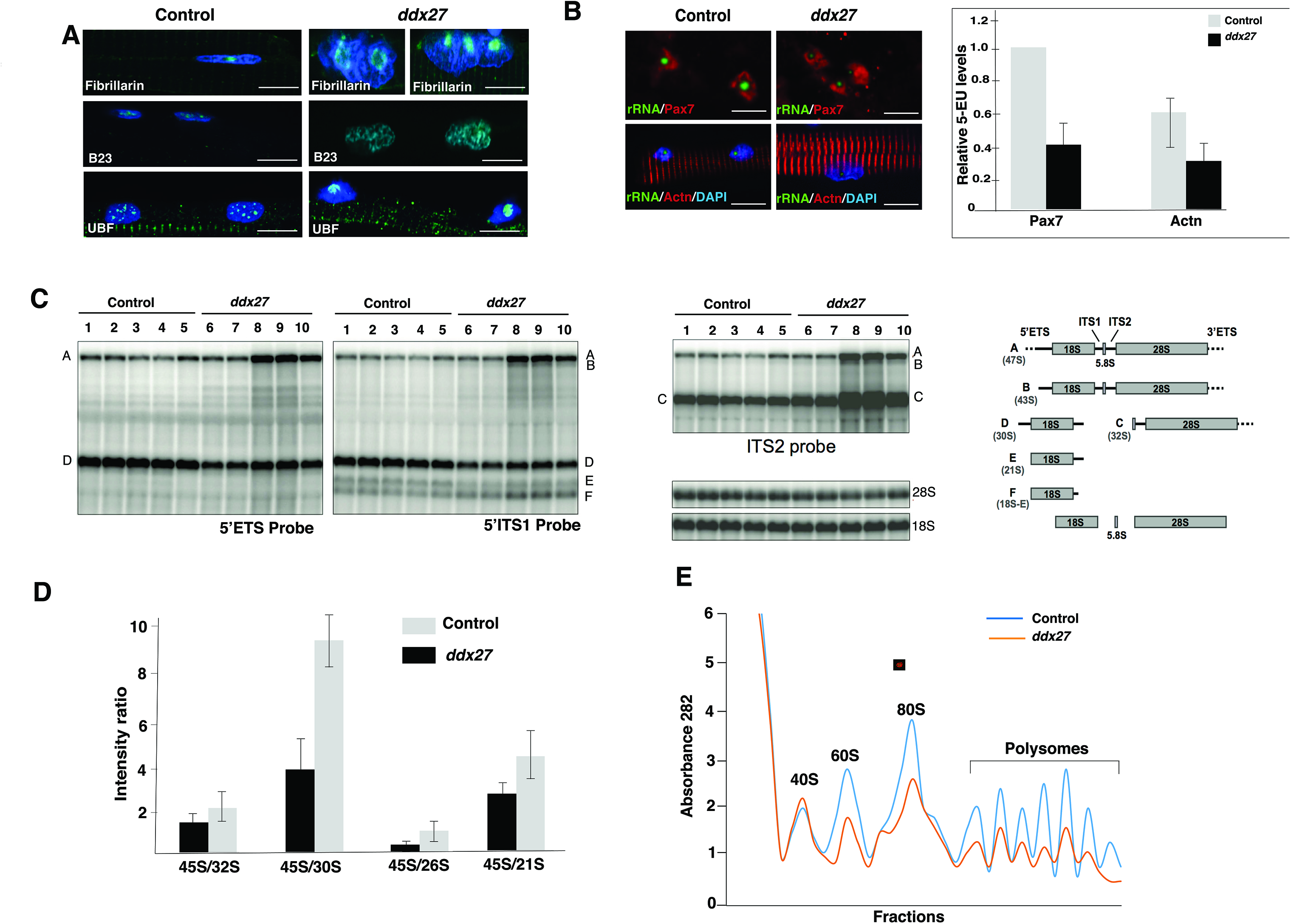
Ddx27 deficiency disrupts nucleolar architecture and rRNA synthesis resulting in ribosomal defects. (A) Immunofluorescence of control and *ddx27* mutant fish with antibodies labeling different nucleolar compartments at 5 dpf (scale bar: 10μm) (B) rRNA transcription was measured in MPCs (labeled with Pax7) or myonuclei (labeled with Actn2/3) at 5 dpf by quantifying the incorporation of 5-ethynl uridine (5-EU). Zebrafish or myofibers were treated with Actinomycin D for two hours to block background transcription and subsequently, were incubated with or without Actinomycin D and freshly synthesized rRNA was quantified by incorporation of 5-EU by fluorescent detection (scale bar: 5μm) (C) Northern blot analysis of total RNAs extracted from skeletal muscles of control and mutant *ddx27* zebrafish larvae (5 dpf). 5’ETS, 5’ITS1 and ITS2 probes were used to identify pre-rRNA and intermediate species targeted different steps of the processing pathways. The pre-rRNA intermediates are described in zebrafish. The corresponding human precursors are indicated into brackets. (D) Quantification of the pre-rRNA intermediates in zebrafish skeletal muscles. (E) Polysomal profiles of skeletal muscle in control and *ddx27* mutant larvae (5 dpf).

Next, to evaluate the role of Ddx27 on pre-rRNA processing, the pre-rRNA maturation pattern was evaluated in the skeletal muscle of *ddx27* mutant zebrafish. Skeletal muscles (containing MPCs and myonuclei) were dissected from control and mutant larval fish (5 dpf) and total RNA was isolated. Northern blotting was performed with different probes that were specific to various precursor rRNAs representing different pre-rRNA processing steps (Fig 5C-D). *ddx27* mutant skeletal muscle displayed significant accumulation of long pre-rRNAs (precursors A and B), corresponding to 47S and 43S pre-rRNAs in the human rRNA processing pathway. Precursors C (corresponding to human 32S pre-rRNA) also accumulated, while precursors to the 18S rRNA (precursors D, E) decreased. This accumulation of early precursors and of precursors C suggests an early defect in the rRNA maturation process and indicative of delayed cleavages in the 5’ETS and 3’ETS. These results are in accordance with recent data showing that depletion of DDX27 in human cells leads to the release of an extended form of the 47S primary transcript [34]. In addition, our data reveal an accumulation of 41S pre-rRNAs and a concomitant decrease of 30S pre-rRNAs, which are indicative of an impaired cleavage at site 2 (Fig 5C).

To study the impact of rRNA maturation defects on ribosomes, we performed ribosomal profiling in control and mutant zebrafish muscle that revealed a significant decrease in free 60S large ribosomal subunits. A reduction of mature 80S monosomes as well as polysomes was also observed in mutant muscles (Fig 5E). Together these studies show that *ddx27* expression is required for the formation of mature rRNA species and thereby for biogenesis of functional polysomes in skeletal muscle.

### Ddx27 deficiency perturbs the translation of specific subset of mRNA repertoires

We next sought to investigate if the ribosomal deficits due to Ddx27 deficiency affect translation of global processes or a specific mRNA repertoire in skeletal muscle. To identify mRNA repertoire exhibiting perturbed translation in Ddx27 deficiency, polysome profiling and subsequent RNA sequencing was performed in *Ddx27* knockout C2C12 myoblasts that exhibit proliferation and differentiation defects similar to zebrafish. Also, Ddx27 is highly expressed in proliferating C2C12 myoblasts suggesting that mRNA species identified from polysomal profiling will potentially be a direct consequence of Ddx27 deficiency. Polysomes (actively translating ribosomes) were purified from control and knockout *Ddx27* C2C12 myoblasts grown in the proliferation media. RNA-sequencing of total and polysome bound mRNA transcripts revealed that 124 transcripts that showed increased and 300 which showed decreased association with polysomes were common in both control and mutant. 1057 mRNA transcripts were exclusively enriched in control polysomes whereas *Ddx27* deficient polysomes showed an enrichment of 286 mRNA transcripts (Fig 6). Data analysis (Fig. S4-5) showed that control polysomes associated transcripts exhibited an enrichment in mRNAs encoding ribosomal, RNA polymerase, RNA degradation and splicing pathways suggesting a high requirement of these RNA metabolic processes during muscle cell growth. On the other hand, in DDX27 deficiency an enrichment of apoptotic and inflammatory pathway genes was observed suggesting that absence of DDX27 activates the atrophic processes in muscle. Interestingly, mutant fish did not exhibit any visible increase in apoptosis. The enrichment of apoptotic transcripts in mutants could potentially be due to an initiation of end stage changes as mutants die by 5 dpf. In addition, mRNAs required for protein biosynthesis (amino acid biosynthesis, aminoacyl-tRNA biosynthesis) showed significantly lower enrichment in mutant polysomes. These result suggest that DDX27 is crucial for the translation of mRNAs that are necessary for generating building blocks for active biosynthesis of proteins and suppression of transcripts associated with atrophic processes during muscle growth.

**Figure 6.**
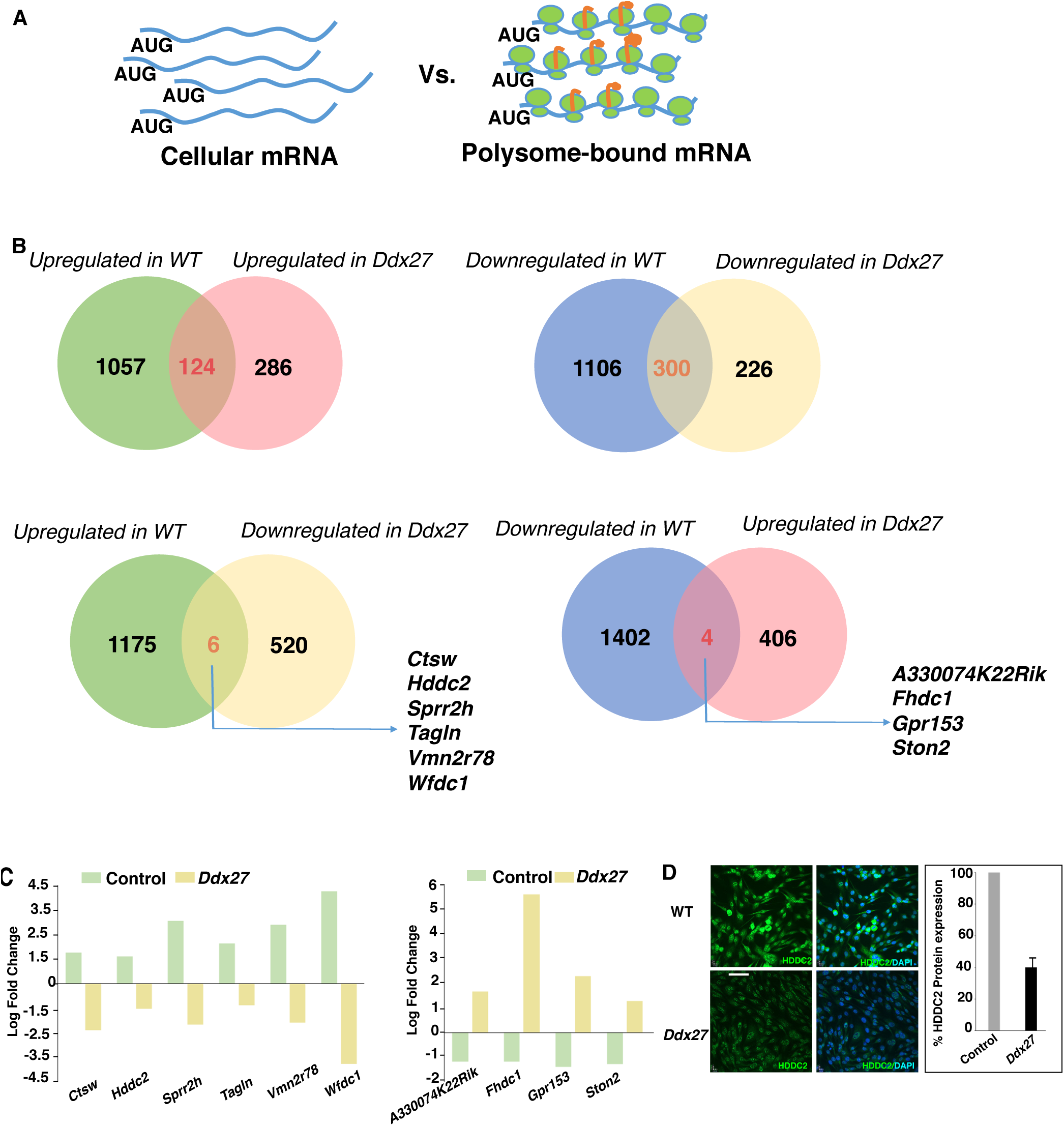
DDX27 is required to regulate specific mRNA repertories in myoblasts. (A) Polysomal profiling and RNA sequencing of total cellular mRNA transcripts and polysomal fractions were performed in control and *Ddx27* mutant C2C12 myoblasts. (B) mRNAs transcripts exhibiting expression values of Log_2_Fold change >+1 were considered upregulated and those displaying Log_2_Fold change <+1 were considered downregulated in the polysomal fraction in comparison to total cellular mRNA. Upregulated and downregulated mRNAs populations in control and *Ddx27* myoblasts were compared (C) Expression of mRNA transcripts exhibiting maximum differences in control and *Ddx27* mutant myoblasts (D) Immunofluorescence and Western blot quantification of HDDC2 expression in control and *Ddx27* proliferating myoblasts (scale bar: 100μm).

Interestingly, a number of signal transduction pathways associated with skeletal muscle growth and diseases are also perturbed in DDX27 deficiency. Mutant polysomes exhibited a reduced association with *Fgfr1* transcripts. FGF-signaling pathway is crucial for skeletal muscle growth and muscle specific ablation of *Fgfr1* impairs proliferation of muscle satellite cells [35]. Additionally, an increase in mRNAs encoding members of MAP Kinase pathway that is associated with precocious differentiation was observed in *ddx27* mutant zebrafish muscles [36]. Lastly, we identified several novel mRNAs that were highly enriched in control polysomes but decreased in DDX27-deficienct polysomes (e.g. *Ctsw, Hddc2, Tagln, Wfdc1 Sprr2h*, and *Vmn2r78*)(Fig 6C). Many of these genes are involved in the maintenance of proliferative state of human ES and ipS cell however, their role/s in myogenesis are not known [37]. To confirmed the validity of our polysomal profiling data and expression of these transcripts in skeletal muscle we analyzed the expression of HDDC2 in control and *Ddx27* mutant myoblasts. Immunofluorescence and Western blotting revealed a significant downregulation of HDDC2 protein in *Ddx27* mutant myoblasts suggesting that HDDC2 may be contributing to DDX27 mediated muscle stem cell defects in skeletal muscle (Fig 6D). A reverse analysis also identified the downregulation of 4 novel transcripts in control myoblasts that were enriched in *Ddx27* polysomes, including *A330074K22Rik, Fhdc1, Gpr153*, and *Ston2* and future studies will be able to identify their functional roles in skeletal muscle. In sum, polysome profiling revealed that DDX27 is required for the translation of mRNAs regulating RNA metabolism and signaling pathway that are crucial for muscle satellite cell proliferation and differentiation. In addition, we identified novel genes that are required for cellular proliferation and future studies on *in vivo* function of these genes in skeletal muscle may help to understand novel processes regulating muscle growth.

### Discussion

In this study, we hypothesized that *in vivo* identification of novel factors regulating myogenesis should help to elucidate the molecular mechanisms that regulate myogenic processes. Indeed, in this work we found that an RNA helicase, Ddx27, regulates skeletal muscle growth and regeneration by controlling ribosome biogenesis and translational processes. Our investigation into mechanisms of Ddx27 function in skeletal muscle growth leads to three major conclusions. First, Ddx27 is required for the skeletal muscle growth by regulating MPCs proliferation and differentiation in to mature myofibers in zebrafish. Second, Ddx27 is critical for skeletal muscle repair of injury in larval fish. Third, DDX27 mediated processes contributing to skeletal muscle growth and repair are primarily regulated at the level of protein translation by ribosomes.

### RNA helicases as regulators of skeletal muscle growth and diseases

DDX27 belongs to the DEAD-box family of RNA helicases which represent a large protein family with 43 members that catalyze the ATP-dependent unwinding of double stranded RNA and variously function in remodeling structures of RNA or RNA/protein complexes, dissociating RNA/protein complexes, or RNA annealing [38]. Studies in yeast and cellular models have shown that several DDX family members regulate different steps of rRNA processing; however, *in vivo* functions of these RNA helicases in vertebrates are still mostly unknown. While this plethora of RNA helicases implies the potential for functional redundancy, it also raises the attractive possibility that RNA helicases might perform a generic, unifying function in neuromuscular system by regulating RNA metabolism. DDX5/p68 RNA helicase promotes the assembly of proteins required for transcription initiation complex and chromatin remodeling during skeletal muscle differentiation [39]. Overexpression of *DDX5* also restores the skeletal muscle function in a mouse model of myotonic dystrophy [40]. RNA helicases (DDX1 and DDX3) play significant roles in muscle diseases by interacting with muscle specific transcription factors or with disease causing genes [41, 42]. We further demonstrate that DDX27 is a nucleolar protein that is highly expressed in muscle progenitor cells. Although nucleolar proteins often have ubiquitous localization, high expression of DDX27 in satellite cells and myoblasts suggests a specialized role for this protein in controlling MPC-regulated processes in skeletal muscle.

### Ddx27 is a regulator of muscle growth and repair in zebrafish

Ddx27 deficiency impairs skeletal muscle growth and regeneration in larval Ddx27-deficient zebrafish. The expression of *pax7* RNA as well as number of Pax7 expressing MPC were found to be significantly reduced in skeletal muscles of *ddx27* fish. This reduced number of Pax7 positive MPC population could be either due to a direct role of Ddx27 in regulating proliferation, premature differentiation or increased cell death. Our studies demonstrate that Ddx27 deficiency leads to a decrease in proliferation of the Pax7-positive muscle progenitor cell population in mutant skeletal muscles by accumulation of cells at the G1 stage of cell cycle (S2H Fig). Considering DDX27 is expressed in murine Pax7 positive cells and proliferating myoblasts (Fig 1), this reduced proliferation of MPC in zebrafish is likely due to a cell autonomous defect. Moreover, no significant increase in apoptosis was observed in *ddx27* mutant muscles suggesting that the reduction in MPC is not a consequence of an increased cell death in mutants. During myoblast differentiation to myotubes, downregulation of Ddx27 expression is associated with an upregulation of MyoD, MyoG and subsequently, myosin heavy chain expression. These data suggest that Ddx27 expression may be required for maintaining the proliferative state of muscle progenitor cells and preventing their differentiation to mature muscles under normal conditions. This is supported by the observation that an absence of Ddx27 results in upregulation of *myoD and myf5* in *ddx27* mutants implicating a precocious differentiation of mutant myoblasts. Previous studies have shown that overexpression of MyoD is sufficient to induce myogenic differentiation suggesting a similar mechanism may be contributing to premature differentiation of Ddx27 deficient MPCs [43, 44].

A lack of muscle repair in mutant larval fish further demonstrated that Ddx27 is also crucial for skeletal muscle regeneration. Skeletal muscle regeneration is a complex process involving migration and proliferation of muscle progenitor cells to the injury site followed by differentiation to form new myofibers or repair the existing damaged myofibers. *In vivo* imaging studies have shown that the process of muscle repair in larval zebrafish is highly similar to that in adult mammalian muscle. In zebrafish, Pax7-marked muscle progenitor cells migrate to the injury site, divide and undergo terminal differentiation and regenerate muscle fibers [9]. Our studies demonstrate an absence of Pax7 expressing MPCs at the injury site in *ddx27* skeletal muscle. This could be either due to a defect in the proliferation of Pax7 muscle progenitor cells as observed by reduced Edu labeling of Pax7 positive nuclei in *ddx27* mutant fish or due to the inability of muscle progenitor cells to migrate to the site of injury and/or fuse with damage fibers. Considering we observed reduced MPCs proliferation as well as a lack of proper myofiber differentiation in Ddx27 deficiency, either or a combination of both of these processes could be contributing to skeletal muscle repair defects in mutant fish. Follow up studies with transgenic reporter lines will help us investigate the contribution of different cell lineages and processes contributing to impaired skeletal muscle regeneration in Ddx27-deficient skeletal muscle.

### Ddx27 regulates nucleolar structure and translational processes crucial for skeletal muscle growth and functions

The nucleolus is a prominent organelle, central to gene expression, in which ribosome synthesis is initiated. Alterations in nucleolar structure are indicators of changes in cellular growth and proliferation, cell cycle regulation and senescence; therefore, identification of mechanisms that guide the nucleolar structure-function relationship *in vivo* is needed. Our studies show that DDX27 is highly expressed in MPCs that regulate skeletal muscle growth and repair and Ddx27 deficiency results in rRNA synthesis defects in the MPC population. A number of recent studies have shown that nucleoli are actively involved in stem cell maintenance [45, 46]. Actively proliferating stem cells and progenitor cells possess large nucleoli that change to smaller foci during differentiation These changes in nucleolar structure are associated with reduced rDNA transcription and ribosome biogenesis during differentiation. Depletion of a number of nucleolar proteins results in reduced cell proliferation, abnormal cell cycle and enhanced differentiation, demonstrating that proper nucleolar function is required for self-renewal of ES cells [47, 48]. During differentiation of mesenchymal progenitors in to osteoblasts, myoblasts or adipocytes, phenotypic regulatory factors critical for each lineage (e.g. Runx2, MyoD, Mgn or C/EBP) suppress rDNA transcription suggesting regulation of rRNA synthesis by cell fate-determining factors is a broadly used mechanism for coordinating cell growth with lineage progression [49]. Notably, we identified that lack of Ddx27 also results in rRNA synthesis defects in myonuclei. As mutant MPCs form mature muscles, rRNA defects are also persisted in differentiated muscle fibers. There are 43 DDX proteins in vertebrates, several of which are also expressed in myonuclei (DDX1, 3 and 5). Therefore, rRNA synthesis defects in *ddx27* mutant myonuclei imply highly specialized roles of Ddx27 in skeletal muscle that are not compensated by other DDX family members. Collectively, these studies suggest that Ddx27 deficiency results in impaired proliferation and differentiation of MPCs leading to defective myofibers. This lack of Ddx27 in MPCs leads to a direct ribosomal defects in MPCs and have an indirect effect on myofibers derived from these MPC population.

Molecularly, we established that Ddx27 regulates rRNA maturation and ribosome biogenesis. DDX27 is highly conserved in evolution, and mutation of the yeast *ddx27* ortholog, *drs1p*, results in 25S rRNA maturation and 60S ribosome subunit biogenesis defects [50]. In zebrafish and mammalian cells, we identified an accumulation of primary transcripts and long-pre-rRNAs reflecting early rRNA maturation defects resulting in a decrease in functional ribosomes. Previous studies have shown that ribosome numbers are increased during skeletal muscle hypertrophy and reduced in atrophied muscles or skeletal muscle diseases [20-22]. However, the regulators of these processes remain to be identified. Our studies demonstrate that DDX27 is required for normal muscle growth by controlling rRNA maturation and ribosomal biogenesis. A number of recent studies have shown that ribosomes of different heterogeneities and functionalities exist and contribute to the translational control of gene regulation by selecting mRNA subsets to be translated under specific growth conditions [5, 51]. For example, analysis of ribosome populations in mouse embryonic stem cells revealed that RPL10 enriched ribosomes preferentially regulate mRNAs controlling cellular growth whereas RPL10 depleted ribosomes exhibited increased binding to mRNA pools regulating stress responses and cell death [52]. Therefore, reduced proliferation of MPCs and terminal differentiation into mature myofibers in zebrafish could potentially be due defects in ribosome biogenesis affecting either the global translation rates or translation of specific subsets of mRNA required for proliferation and differentiation of MPC population. Polysomal profiling identified reduced association of mRNAs in FGF and MAPK signaling pathways that are known to regulate different stages of skeletal muscle growth in both cell autonomous and non-autonomous manners. These signaling pathways are associated with activation and myogenic commitment of muscle stem cells and thus could be contributing to the skeletal muscle defects observed in Ddx27 deficiency in zebrafish [35, 36]. Polysomal profiling also revealed an enrichment of transcripts regulating RNA metabolism pathways in control polysomes that was lacking in the mutant muscles. Defects in RNA metabolism underlie disease pathophysiology in a number of neuromuscular diseases [53]. Therefore, identification of critical regulators of RNA based processes that contribute towards myogenesis is essential in order to understand the molecular basis of pathological changes in disease conditions. Interestingly, many pathways altered in mutant myoblast are also crucial for myofiber hypertrophy. Identification of these different processes in control and mutant muscles signifies an additional regulatory layer of translational regulation controlled by DDX27 that fine tunes the crucial processes associated with skeletal muscle growth. Interestingly, most of the translational changes observed in both control and mutant skeletal muscle are associated with genes that were previously not known to play significant roles in myogenesis. In particular, dysregulation of genes associated with cellular pluripotency and membrane remodeling suggests novel roles for these factors in skeletal muscle biology. Future work targeting these genes will help to illuminate additional processes that are crucial for myogenesis. This work provides new insight into nucleolar function in skeletal muscle growth, and opens a new avenue to explore the specific roles of nucleolar proteins and ribosome biogenesis in normal and disease muscle.

## EXPERIMENTAL PROCEDURES

### Zebrafish lines

Fish were bred and maintained using standard methods as described (56). All procedures were approved by the Boston Children’s Hospital Animal Care and Use Committee. Wild-type embryos were obtained from Oregon AB line and were staged by hours (h) or days (d) post fertilization at 28.5°C. Zebrafish embryonic (0-2 days post fertilization) and larval stages (3-5 dpf) have been defined as described previously [54].

### Skeletal muscle regeneration

Cardiotoxin-induced muscle regeneration studies in zebrafish were performed following previously published protocols [55]. Briefly, control and *ddx27* mutant larvae (3 dpf) were anesthetized and immobilized by embedding into 3% low melting agarose. Cardiotoxin (10mM, 1 μl) was injected into dorsal somite muscles, and fish (8-10, in 4 independent experiments) were analyzed at 5 dpf by immunofluorescence analysis. Muscle degeneration and regeneration in mice was performed as described previously [56].

### Skeletal Muscle Functional Analysis

Functional experiments were performed as previously described [57]. Briefly, fish were studied in a bicarbonate buffer of the following composition: (in mM) 117.2 NaCl, 4.7 KCl, 1.2 MgCl_2_, 1.2 KH_2_PO_4_, 2.5 CaCl_2_, 25.2 NaHCO_3_, 11.1 glucose (Dou et al., 2008). 4-5 dpf larvae were anesthetized in fish buffer containing 0.02% tricaine and decapitated. The head tissue was used for genotyping. The larval body was transferred to a small chamber containing fish buffer equilibrated with 95% O_2_, 5% CO_2_ and maintained at 25 ^°^C. The larval body was attached to an isometric force transducer (Aurora Scientific, Aurora, Ontario, CAN, model 403A) and position motor (Aurora Scientific model 308B) using a 10-0 monofilament tie placed at the gastrointestinal opening and another tie attached several myotomes proximal from the tip of the tail. Twitches (200 μs pulse duration) and tetani (300 Hz) were elicited using supramaximal current delivered to platinum electrodes flanking the preparation. All data were collected at the optimal preparation length (L_o_) for tetanic force. At the conclusion of the experiment, images of the preparations width and depth at L_o_ were obtained by carefully rotating the preparation about the gastrointestinal opening attachment point. Each image was analyzed by ImageJ using an internal length calibration. Preparation cross-sectional area (CSA) was calculated from width and depth measurements assuming the preparations cross-section was elliptical. Forces were calculated as active force, i.e. peak force minus the unstimulated baseline force, and are presented in absolute terms (valid because all larvae were attached at a consistent anatomical landmark, the gastrointestinal opening) as well as normalized to preparation CSA. The maximal rate of twitch tension development was determined as the maximal derivative of the force by time response between the onset of contraction and peak force. Likewise, the maximal rate of twitch tension relaxation was calculated as the first derivative of the force by time relaxation response, ranging from peak force until force had declined to approximately baseline. Statistical differences in the *ddx27* (n = 12) and control (n = 10) group means were evaluated by a two-sample t-test.

### Single Myofiber culture and immunofluorescence

Whole mount immunofluorescence in zebrafish was performed as described previously [58]. For immunofluorescence studies in zebrafish myofiber culture, previously published protocol was followed and 30-40 myofibers were analyzed in each condition [59]. Paraformaldehyde (4%) or methanol (100%) were used as fixative for different antibodies. EDL culture and immunofluorescence was performed using previously published studies [60]. Primary antibodies used in this study were: a-actinin (1:100; Sigma, A7811), RYR1 (1:100; Sigma, R-129), Fibrillarin (1:50; Santa Cruz, sc-25397), B23 (1:50; Santa Cruz, sc-5564), UBF (Sigma, 1:50, HPA006385), DDX27 (1:50, Santa Cruz, sc-81074), Pax7 (1:20; Developmental Studies Hybridoma Bank), Mef2 (1:20, Santa Cruz, sc-17785). For phalloidin staining, paraformaldehyde fixed embryos/larvae were incubated with phalloidin (1:40, Thermo Fisher Scientific, A12379) (and with primary antibody; for double immunofluorescence), overnight at 4°C followed by incubation with secondary antibody (if primary antibody was used). Nuclear staining was done using DAPI (Biolegend, 422801). Secondary antibodies (Thermo Fisher Scientific) were used between 1:100-1:250 dilutions.

### C2C12 myoblasts culture and differentiation

Mouse C2c12 cells were cultured in growth medium consisting of DMEM supplemented with 20% fetal bovine serum. To induce differentiation, growth medium was replaced with the differentiation medium consisting of DMEM supplemented with 2% horse serum. Cells were maintained in the differentiation medium for 5 days. Western blot analysis was performed with Ddx27(1:100, Santa Cruz, sc-81074), MyoD (1:200, Santa Cruz, sc-760), MyoG (1:200, DHSB, F5D) and MF20 (1:50, DHSB).

### rRNA transcription, maturation and ribosome analysis

rRNA transcription was detected using the Click-iT RNA Alexa Fluor imaging kit (C10329, Invitrogen) as described previously [61]. Briefly, to detect the synthesis of rRNAs in the nucleoplasm, control or *ddx27* knockout zebrafish or cultured myofibers (5 dpf) were treated with 1μg/mL actinomycin for 20 minutes and then incubated for 2 hours with 1mM 5-ethyl uridine (5-EU) in the presence or absence of actinomycin, fixed with paraformaldehyde (4%) and incubated with Click-iT reaction cocktail for 1 hour. This was followed by immunofluorescence with Pax7 in zebrafish (1:20, DHSB) to detect MPCs and a-actinin (1:250, Sigma: A7811) in cultured fish myofibers to label myonuclei. DNA was counterstained with DAPI. The average 5-EU signal intensity in the nucleoplasm was measured from 20 nuclei per experiment using the ImageJ program. The level of rRNA was determined by actinomycin-treated and non-treated nuclei in control and mutant fish or myofibers.

In order to analyze the precursors to the 28S and 18S rRNAs, total RNA samples were isolated from skeletal muscle of control and mutant larvae (n=100) and separated on a 1.2% agarose gel containing 1.2% formaldehyde and 1× Tri/Tri buffer (30 mM triethanolamine, 30 mM tricine, pH 7.9). RNAs were transferred to Hybond N^+^ nylon membrane (GE Healthcare, Orsay, France) and cross-linked under UV light. Membrane hybridization with radiolabeled oligonucleotide probes was performed as described (Preti, 2013). Signals were acquired with a Typhoon Trio PhosphorImager and quantified using the MultiGauge software. The human probes were: 5′-TTTACTTCCTCTAGATAGTCAAGTTCGACC-3′ (18S), 5′-CCTCGCCCTCCGGGCTCCGTTAATGATC-3′ (5′ITS1), a mixture of 5′-CTGCGAGGGAACCCCCAGCCGCGCA-3′ (ITS2-1) and 5′-GCGCGACGGCGGACGACACCGCGGCGTC-3′ (ITS2-2), 5′-CCCGTTCCCTTGGCTGTGGTTTCGCTAGATA-3′ (28S), 5′-GCACGCGCGCGCGGACAAACCCTTG-3′ (28S-3′ETS). The zebrafish probes were: 5′-GAGGGAGGCGCGTCGACCTTCGCTGGGC-3′ (3′ETS), 5′-CAGCTTTGCAACCATACTCCCCCCGGAAC-3′ (18S), 5′-GAGATCCCCTCTCGAACCCGTAATGAT-3′ (ITS1), 5′-GAGCGCTGGCCTCGGAGATCGCTGGGTCGC-3′ (ITS2), 5′-CCTCTCGTACTGAGCAGGATTACTATTGC-3′ (28S). For ribosome profile analysis zebrafish (100 larvae) were treated with cycloheximide (Sigma, 100ug/ML) for 10 minutes at room temperature. Subsequently, skeletal muscle was dissected and flash frozen in liquid nitrogen in lysis buffer (10mM Tris-Cl, pH 7.4, 5mM MgCl2, 100mM KCl, 1% TritonX-100). To purify ribosomal fractions, cell lysate (0.75 ml) was layered on 10-50% sucrose gradient and centrifuged at 36,000 rpm using SW-41 Ti rotor, 2 hours at 4^°^C. The fractions were collected and were analyzed at 254nm.

### Polysomal profiling

Polysome ribosome fractions were prepared from control and *Ddx27* C2C12 myoblasts. Equal number of control and Ddx27 knockout C2C12 myoblasts were plated in 10 cm dishes in the proliferation media and collected after 24 hours for polysomal analysis. Polysomes were isolated as described above for zebrafish muscle and fractions were pooled together, treated with proteinase K and RNA was isolated using acid phenol-chloroform extraction and ethanol precipitation. As Ddx27 myoblasts exhibit a reduced proliferation, equal amounts of proteins were used to fractionate polysomes from control and KO myoblasts. Deep sequencing libraries were generated and sequenced as described [62]. Ribosomal profiling was repeated in triplicate and principal component analysis was performed to identify variation between samples. The Spearman R^2^ was > 0.9 for all replicates (except one sample) that was subsequently removed from follow up analysis. Splice-Aware alignment program STAR was used to map sequencing reads to *Mus musculus* (mm10 build). R package “edgeR” was employed to identify differential gene expression calls from these sequence reads. Gene expression was considered to be up-regulated if log2FC> +1 or downregulated if the log2FC< −1 (FC=fold change of average CPM) with respect to the condition being compared at a false discovery rate <0.05. Functional ontological classification of different gene lists was performed by DAVID.

### Proliferation and apoptosis Assays

To analyze the proliferating cells in C2C12 cell cultures and zebrafish embryos, Edu labeling was performed. For zebrafish, embryos (48 hrs) were placed in 1mL of 500μM EdU/10% DMSO in E3 media and incubated on ice for 2 h. Embryos were transferred back to the incubator at 28.5 °C and samples were collected at desired time intervals and fixed in methanol (-20°C, 20 mins). Fish were permeabilized with 1% tritonX-100/PBS for 1 hr and Click-iT reaction cocktail (Thermo Fisher Scientific) was added and incubated in dark for 1 hr at room temperature. Immunofluorescence was performed with Pax7 antibody (1:10 DHSB) and analyzed (10-12 larvae) by confocal microscopy. For analyzing proliferating C2C12 cells, equal number of control and mutant cells (to 40-50% confluency) were plated for 12-14 hours. 2X EdU solution was added to the cells and incubated for 2 hrs at 37°C. After incubation cells were permeabilized with 0.5% triton-100 and labeled with Click-iT reaction cocktail as described for zebrafish. Immunofluorescence was performed with Pax7 antibody (DHSB) and nuclei were stained with DAPI. Apoptosis was performed on 3-4dpf zebrafish larvae by *in situ* cell detection kit (Roche) or western blot analysis using caspase 3 antibody. (ab13847, Abcam)

### Zebrafish locomotion assay

Zebrafish swimming behavior was quantified by an infra-red tracking activity monitoring system (DanioVision, Noldus, Leesburg, VA, USA). Control or *ddx27* mutant larvae were placed individually into each well of a 24 well plate in dark for 10 minutes. The activity of these larvae was recorded during a follow-up light exposure of 20 minutes. Four independent blind trials were performed and mean velocity, total distance and cumulative duration of movement were recorded. Reported values reflect an average of 30-35 control or mutant larval fish.

### Quantification and statistical analysis

Quantification of myofiber size, area, western blots and northern blots were performed using the ImageJ program. Data were statistically analyzed by parametric Student *t*-test (two tailed) and were considered significant when P<0.05. All data analyses were performed using XLSTAT software. Additional detailed experimental details are provided in the supplemental material.

## Acknowledgements

The authors are grateful to Nathalie Montel-Lehry for technical support with rRNA analyses.

